# CYB-4/CDK-1 define temporal windows for chromosome morphogenesis during meiotic prophase in *C. elegans*

**DOI:** 10.64898/2026.10.06.756462

**Authors:** Amirhossein Pezeshki, Steven J.M. Jones, Monique Zetka

## Abstract

Meiotic chromosome segregation requires the earlier coordination of chromosome morphogenesis with cell-cycle progression. In *C. elegans,* chromosome remodelling in preparation for the meiotic divisions is regulated by the master cell-cycle regulator CDK-1, but the cyclin directing its activity in prophase was unknown. Here, we identify CYB-4 as a meiosis-specific B-type cyclin that controls the timing of chromosome axis assembly and bivalent restructuring. A genetic suppressor analysis using *him-3(vvc)* hypomorphs defective in chromosome axis assembly revealed a role for CYB-4 at two key prophase transitions. Loss of CYB-4 delays axis formation, resulting in synapsis defects and the extension of CHK-2 activity, indicating that it is required for timely axis assembly. While recombination progression is also delayed in *cyb-4* mutants, it ultimately recovers to form wild-type levels of crossovers (CO). Despite the presence of COs at late pachytene, however, the bivalent remodeling and chromosome compaction that are hallmarks of pachytene exit are also delayed, indicating that loss of CYB-4 delays a second cell-cycle transition following CO formation. These defects are ultimately resolved before prophase exit and *cyb-4* mutants show wild-type-like levels of fertility, indicating that CYB-4 is required for the scheduling of meiotic events rather than if they occur. Together, our findings provide evidence that CYB-4 directs CDK-1 activity during meiotic prophase to regulate cell cycle-transitions featuring large-scale changes to chromosome structure. We propose that temporal constraints implemented by CYB-4/CDK-1 are essential for the sharp transitions at prophase entry and pachytene exit that coordinate chromosome morphogenesis with meiotic progression.

## Introduction

Meiosis is a specialized cell division in which a single round of DNA replication is followed by two consecutive rounds of chromosome segregation to generate haploid gametes. A defining feature of meiosis I is the extended prophase that is marked by chromosome pairing, homologous recombination (HR), and crossover formation (COs) (Zickler and Kleckner 2023; Ishiguro 2024). Following meiotic entry, chromosomes are reorganized into arrays of chromatin loops that are anchored to a proteinaceous chromosome axis (Ur and Corbett 2021). This higher-order chromosome architecture is required for homolog pairing and provides a structural platform for subsequent meiotic events, including synapsis and recombination (Grey and de Massy 2021). During synapsis, pairing between homologs is stabilized by the synaptonemal complex (SC), a zipper-like proteinaceous structure that is also required for CO formation (Gordon and Rog 2023). Meiotic recombination is initiated by the formation of programmed DNA double-strand breaks (DSBs) by the conserved topoisomerase-like SPO-11 (Harper et al. 2025). Although most DSBs are repaired as noncrossovers, a subset mature into COs that can direct chromosome segregation at the first meiotic division (Zickler and Kleckner 2015).

The chromosome axis is composed of sister chromatid cohesin complexes (SCC) and meiotic HORMA domain-containing proteins (known as mHORMADs) (reviewed in Gu et al. 2022). mHORMADs were originally identified through sequence homology to the budding yeast protein <u>Ho</u>p1 and the checkpoint proteins <u>R</u>ev7 and <u>Mad</u>2 (Aravind and Koonin 1998). Structural studies of Mad2 revealed that the HORMA domain can transition between structurally distinct open and closed conformations during interactions with its binding partners (Luo and Yu 2008). In *C. elegans*, the ordered recruitment of four mHORMAD proteins to chromosomes at meiotic entry is required to assemble the meiotic chromosome axis: HTP-3, HIM-3, and the highly related paralogs HTP-1 and HTP-2 (referred to as HTP-1/2) (Couteau et al. 2004; Couteau and Zetka 2011; Martinez-Perez and Villeneuve 2005; Goodyer et al. 2008; Nabeshima et al. 2004). HTP-3 localization is cohesin dependent, and it subsequently serves as a scaffold for axis assembly through binding of HIM-3 and HTP-1/2 to closure motifs located within its disordered C terminal region (Kim et al. 2014). Despite their physical proximity at the axis, the mHORMADs perform distinct functions in processes that are tightly coordinated. HTP-3 is essential for chromosome axis assembly and DSB induction (Goodyer et al. 2008; Severson et al. 2009); HIM-3 is required for homolog pairing, SC assembly, and the inter-homolog recombination bias (Zetka et al. 1999; Couteau et al. 2004; Sato-Carlton et al. 2020); and HTP-1/2 cooperate in regulating synapsis (Couteau and Zetka 2005; Martinez-Perez and Villeneuve 2005; Nabeshima et al. 2004; Silva et al. 2014). Together, these proteins establish a chromosome architecture that supports CO formation and chromosome segregation. In fungi, plants, and mammals, the chromosomal association of mHORMADs is regulated by PCH2/TRIP13, a member of the AAA+ ATPase family (reviewed in Bhalla 2023). In the case of *C. elegans*, however, mHORMAD localization is independent of PCH-2, indicating that distinct regulatory mechanisms govern chromosome axis assembly in this organism (Bhalla 2023). Recent evidence has revealed that a nuclear <u>c</u>haperonin-<u>c</u>ontaining <u>T</u>CP-1 (CCT/TRiC) complex is required for timely assembly of mHORMADs into the chromosome axis, revealing a regulatory function for protein folding in meiotic chromosome morphogenesis (Zetka et al. 2025).

Progression through the cell cycle is coordinated by cyclin-dependent kinases (CDKs) and their cyclin partners. Cyclins confer substrate specificity and spatiotemporal regulation to CDKs, enabling phosphorylation of target proteins that drive cell-cycle transitions (Pellarin et al. 2025). In *C. elegans*, CDK-1 is required for both mitotic and meiotic M-phase entry and partners with three B-type cyclins (reviewed in Kipreos and van den Heuvel 2019); CYB-1 and CYB-3 are essential cyclins required for both mitosis and meiosis, while CYB-2.1 and its near-identical paralog CYB-2.2 are largely redundant and loss of both causes only minor developmental defects (van der Voet et al. 2009; Rabilotta et al. 2015; Yoon et al. 2017; Pellarin et al. 2025). CDK-1 is required for MSP-dependent oocyte maturation and promoting nuclear envelope (NE) breakdown prior to fertilization (Boxem et al. 1999; McCarter et al. 1999; Miller et al. 2001; Burrows et al. 2006; reviewed in Huelgas-Morales and Greenstein 2018). During meiotic prophase, CDK-1-mediated phosphorylation of the SC component SYP-1 generates a docking site for the Polo-like kinase PLK-2 (Brandt et al. 2020). PLK-2 subsequently directs chromosome remodeling to establish long and short chromosome arms of the bivalent relative to CO sites at pachytene exit (Harper et al. 2011; Labella et al. 2011; Brandt and Kim 2021). CDK-1 is also required for the recruitment of AIR-2/Aurora B to the short arm, thereby coupling REC-8 phosphorylation to oocyte maturation and contributing to the stepwise release of sister chromatid cohesion during the meiotic divisions (Ferrandiz *et al*. 2018). Despite its roles at multiple steps of meiotic prophase, the functions of the known cyclin partners (CYB-1, CYB-2.1/2.2, and CYB-3) of CDK-1 are largely confined to the oocyte meiotic divisions and early embryonic mitoses (van der Voet *et al*. 2009; Deyter *et al*. 2010; Michael 2016), indicating that another cyclin directs CDK-1 activity during early meiosis. Here, we identify CYB-4 as a previously uncharacterized meiosis-specific B-type cyclin that directs CDK-1 activity to coordinate chromosome morphogenesis with pathways mediating chromosome pairing, CO formation, and bivalent remodeling during prophase progression.

## Materials and methods

### *C. elegans* strains and conditions

*C. elegans* strains were maintained at 20°C on nematode growth medium (NGM) plates seeded with OP50 *E. coli* bacteria (Brenner 1974). The N2 Bristol strain was used as the wild-type reference, and all mutant strains were constructed in this background. Strains used in this study are summarized in Supplementary Table 1.

### Scoring embryonic lethality, incidence of males and number of DAPI bodies in diakinesis nuclei

To score embryonic lethality (Emb) and incidence of males (Him) phenotypes 10 L4 larvae were transferred to 10 NGM plates at 20°C to lay eggs. The worms were transferred to fresh plates every 12h for 3 days and the number of eggs laid was scored followed by scoring of surviving adult hermaphrodites and male progeny. To compare Emb and Him among genotypes a chi-square test was used and P < 0.05 considered significant (Zetka et al. 2025).

To quantify DAPI body numbers in diakinesis nuclei, worms were dissected and the germlines were stained with DAPI (4′,6-diamidino-2-phenylindole) to visualize meiotic chromosomes. Dissected germlines were imaged, and z-stacked projections were used to count DAPI bodies at -1 and -2 diakinesis oocytes at the proximal end of the germline (Zetka et al. 2025). For each genotype a minimum of 40 nuclei were analyzed. To compare DAPI body numbers among genotypes a Kruskal-Wallis test was used and P < 0.05 considered significant.

### Germline dissection, antibody staining and imaging

To study *C. elegans* germlines 20-30 L4 larvae were placed on plates at 20°C and dissected after 24h. Dissections were performed in 25μl of 1X PBS on hydrophobic slides. Worms were cut near the head using needles, allowing germlines to extrude due to internal pressure. The excess PBS was removed and the germlines were fixed by adding 15μl 1% PFA (paraformaldehyde) in 1X PBS for 5 minutes. Slides were freeze-cracked in liquid nitrogen and were put in 99% methanol at -20°C for 3-5 minutes. Samples were washed three times for 15 min each in 1X PBS containing Tween-20 (PBS-T) (BioShop Canada Inc, Burlington, ON). Samples were blocked using 1% BSA (bovine serum albumin) in PBS-T for 40-60 minutes in a dark box at room temperature. Samples were incubated overnight with primary antibodies diluted in 1% BSA in PBS-T. Following primary antibody incubation, slides were washed three times for 15 min each in PBS-T. Samples were incubated with secondary antibodies diluted in BSA in PBS-T for 2h at 4°C in a dark box. Slides were washed three times for 15 min each in PBS-T. Ten microliters of DAPI solution diluted in VECTASHIELD antifade mounting medium (Vector Laboratories, Burlingame, CA, USA) were added to each slide and a coverslip was placed over the samples and sealed with nail polish. Prepared slides were imaged using a wide-field microscope (Zeiss Axio Observer inverted microscope platform). For each genotype at least 20 germlines were analyzed and representative images were acquired. Images were processed using Zeiss Zen (blue edition, version 3.7.97.04000) to make maximized or partial Z projected 2D images and were subsequently deconvolved. Final figures were prepared using Adobe Photoshop (version 25.12.0).

The following primary antibodies were used in this study: rabbit α-HIM-3 1:200 (Couteau et al. 2004), guinea pig α-HTP-3 1:700, rabbit α-HTP-3 1:200 (Goodyer et al. 2008), rabbit α-HTP-1/2 1:300 (Couteau and Zetka 2005), rabbit α-PLK-2 1:100 (Labella et al. 2011), rabbit α-RAD-51 1:200, rabbit α-RAD-51 1:2000 (Novus Biologicals, #29480002), guinea pig α-SYP-1 1:800, guinea pig α-pSUN-1 (against serine 8) 1:1300 (Woglar et al. 2013), rabbit α-SYP-1 1:1000, rabbit α-GFP 1:2000 (Abcam, ab1218), rabbit α-COH-3/4 1:200 (Severson and Meyer 2014), mouse α-GFP 1:200 (Abcam, ab291), mouse α-FLAG 1:500 (Sigma Aldrich F1804), monoclonal mouse anti-HA 1:1000 (Cell Signaling, #2367), monoclonal rat α-HA HRP-conjugated high affinity 1:2000 (Roche, #11867423001), monoclonal mouse anti-FLAG HRP-conjugated 1:5000 (Sigma, #A8592), polyclonal chicken α-GFP 1:5000 (AbCam, #ab13970), polyclonal rabbit α-Histone H3 1:100,000 (AbCam, #ab1791). And the secondary antibodies used in this project are as follow: Alexa Fluor 488 goat α-rabbit IgG 1:1000 (Invitrogen, A11034), Alexa Fluor 488 Donkey α-guinea pig IgG 1:600 (Jackson ImmunoResearch #166451), Alexa Fluor 488 goat α-guinea pig IgG 1:1000 (Invitrogen A-11073), Cy3 donkey α-rat IgG 1:300 (Jackson ImmunoResearch #55721), Alexa Fluor 555 goat α-rabbit IgG 1:1000 (Invitrogen, #A21429), Alexa Fluor 555 goat α-guinea pig IgG 1:1000 (Thermo Fisher A-21435), Alexa Fluor 488 donkey α-mouse IgG 1:500 (Jackson ImmunoResearch #106498).

### Defining prophase stages and colocalization coefficient measurements

To define stages of meiotic prophase I, the TZ was identified by the presence of polarized, crescent-shaped nuclei and/or by staining with meiosis-specific markers such as HIM-3, the kleisin COH-3/4, and phosphorylated SUN-1 in nuclear aggregates. The meiotic entry region was defined as the region where there is a mixture of premeiotic nonpolarized nuclei and meiotic nuclei as defined by nuclear polarization and localization of meiosis-specific COH-3/4 (Jantsch et al. 2007; Severson and Meyer 2014). The end of the meiotic entry region was defined as the first row of polarized and COH-3/4 positive nuclei. The extent of the meiotic entry region was measured and normalized to germline length, defined as the distance from the distal tip to the germline bend. Early pachytene (pachytene entry) was defined as the point at which ≥50% of nuclei were no longer polarized, and pachytene exit was defined as the first nuclei that followed the last nuclei showing synapsed chromosomes. Mid-pachytene was defined as the middle third of the region between early pachytene and pachytene exit (Zetka et al. 2025).

Protein localization was quantified by measuring the co-localization coefficient (CoC) between the protein of interest and a reference protein. CoC values were obtained using the co-localization analysis tool in ZEISS ZEN Microscopy Software (RRID:SCR_013672), version 3.7. Images were deconvolved and subsequently Z-projected prior to analysis and CoC was measured after the background threshold was determined by analyzing a portion of the gonad containing no visible signal for both channels. Regions of interest were defined by drawing a circle around each nucleus, and the co-localization between channels was quantified using Pearson’s correlation coefficient. To compare CoC among genotypes a Kruskal-Wallis test followed by Dunn’s pairwise comparison test was used and P < 0.05 considered significant (Zetka et al. 2025).

### CRISPR-CasG directed mutagenesis

Mutations of interest were generated using CRISPR–Cas9-mediated genome editing (Kim and Colaiácovo 2019) by microinjecting CRISPR reagents into 24 h post-L4 young adult N2 wild-type animals, except for EZ598 strain, which was generated in CB4856 Hawaiian background. The injection mix consisted of Cas9 protein (2 mg/mL; PNA Bio Inc), tracrRNA (8 µg/µL), *dpy-10* crRNA (8 µg/µL), *dpy-10* repair template (500 ng/µL), gene-specific crRNA (8 µg/µL), and a gene-specific repair template (1 µg/µL), along with KCl (1 M) and HEPES (200 mM, pH 7.4). F1 progeny were screened for the desired mutation by PCR followed by restriction fragment analysis and confirmed by Sanger sequencing. The homozygous mutants containing the mutation of interest were outcrossed three times with N2 wild-type strain to remove background mutations. The guide RNA, DNA template, primer sequences are described in Supplementary Table 2.

## Results

### Identification of CYB-4 as a suppressor of chromosome axis defects in *him-3(vvC)* mutants

*him-3(vvc)* hypomorphic mutants exhibit delayed HIM-3 axis localization, resulting in defective homolog pairing, delayed SC assembly, and loss of CO formation (Couteau and Zetka 2005; Zetka et al. 2025). Suppression of the low fertility of *him-3(vvc)* mutants was used to identify genes that regulate chromosome axis morphogenesis (Zetka et al. 2025), and here we describe the characterization of the *vv50* mutant. To identify the mutation corresponding to *vv50*, we combined <u>si</u>ngle-<u>n</u>ucleotide <u>p</u>olymorphism (SNP) mapping with <u>w</u>hole-<u>g</u>enome <u>s</u>equencing (WGS) (Davis et al. 2005; Smith et al. 2016) and found a G-to-A substitution in the second exon of the uncharacterized gene *T12CS.7* that introduced a premature stop codon at glutamine 52 (Q52*) (Fig. 1a). To confirm that this mutation was responsible for the suppression of *him-3(vvc)*, *Ǫ52\** was introduced into T12C9.7 in the wild-type background using CRISPR-Cas9 genome editing (Kim and Colaiácovo 2019). The engineered *T12CS.7(vv17c)* allele suppressed *him-3(vvc)* phenotypes to the same extent as the original *vv50* isolate; embryonic lethality in *him-3(vvc)* mutants was reduced from 86% to ∼42% in the presence of either *vv50* or *vv17c* (Fig. 1a and b). In wild-type germlines, the diakinesis nuclei invariably contain 6 DAPI-bodies representing 6 paired chromosomes joined by a single chiasma. While *him-3(vvc)* mutants contained an average of 10.2 DAPI-stained bodies at diakinesis, indicative of defective CO formation, *vv50; him-3(vvc)* and *T12CS.7(vv17c*); *him-3(vvc)* mutants both resulted in decreases to an average of ∼8 DAPI bodies (Fig. 1c). Together, these results demonstrate that the *T12CS.7(vv17c*) mutation fully recapitulates the *vv50* suppression phenotype, and confirm that *T12CS.7* is the causal gene.

**Fig. 1.**
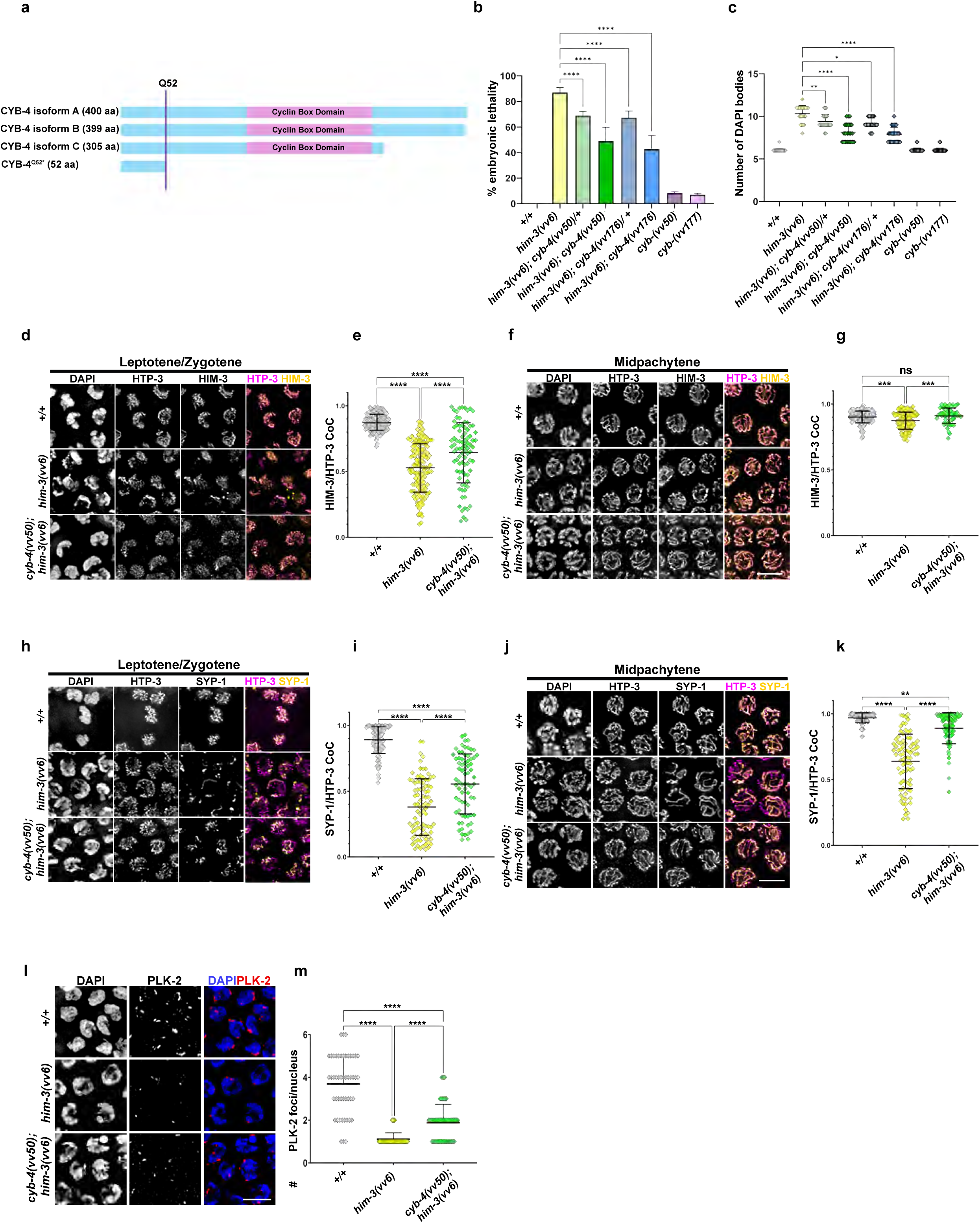
Delayed axis assembly, SC formation, defective CO formation and chromosome segregation in *him-3(vvC)* mutants are partially rescued by *cyb-4(vv50)*. a) Schematic diagram showing CYB-4 protein isoforms and truncated CYB-4*^vv50^*. b) Bar graphs showing % of embryonic lethality in the indicated genotypes. A chi-square test was used for statistical analysis and P < 0.05 was considered significant (n = 10 for all genotypes). c) Scatterplots showing DAPI body counts in diakinesis nuclei for the indicated genotypes (a minimum of 43 nuclei were counted for each genotype). d) Immunofluorescence micrographs showing TZ nuclei stained for DNA, HTP-3 and HIM-3. e) Scatterplot showing colocalization coefficient (CoC) of HIM-3/HTP-3 in the TZ nuclei of the indicated genotypes (n for wild type = 147, n for *him-3(vvc)* = 145 and n for *him-3(vvc); cyb-4(vv50)* = 85). f) Immunofluorescence micrographs of midpachytene nuclei stained for DNA, HTP-3 and HIM-3. g) Scatterplot showing colocalization coefficient (CoC) of HIM-3/HTP-3 in the midpachytene of the indicated genotypes (n for wild type = 124, n for *him-3(vvc)* = 124 and n for *him-3(vvc); cyb- 4(vv50)* = 56). h) Immunofluorescence micrographs showing TZ nuclei stained for DNA, HTP-3 and SYP-1. i) Scatterplot showing the CoC of SYP-1/HTP-3 in the TZ of the indicated genotypes (n for wildtype = 98, n for *him-3(vvc)* = 87 and n for *him-3(vvc); cyb-4(vv50)* = 72). j) Immunofluorescence micrographs showing midpachytene nuclei of stained for DNA, HTP-3 and SYP-1. k) Scatterplot showing the CoC of SYP-1/HTP-3 in the midpachytene nuclei of the indicated genotypes (n for wildtype = 82, n for *him-3(vvc)* = 82 and n for *him-3(vvc); cyb-4(vv50)* = 71). l) Immunofluorescence micrographs showing nuclei of the TZ stained for DNA and PLK-2. m) Scatterplot showing the number of PLK-2 foci per nucleus in the indicated genotypes (n for wildtype = 49, n for *him-3(vvc)* = 49 and n for *him-3(vvc); cyb-4(vv50)* = 61). Kruskal-Wallis test was used for statistical analysis unless otherwise indicated and P < 0.05 was considered significant. ns: not significant, *: P < 0.1, **: P < 0.01, ***: P < 0.001, ****: P < 0.0001. Scale bars, 5µm.

Analysis of the predicted protein sequence of *T12CS.7* identified two conserved cyclin-box domains (Supplementary Fig. 1a) (Blum et al. 2025); orthology analysis placed T12C9.7 within the B-type cyclin family (Supplementary Fig. 1b) (Ruan et al. 2008), and its alignment with human and *C. elegans* cyclins revealed that *T12CS.7* clusters most closely with cyclin B3 proteins from both species (Supplementary Fig. 1c and d). Collectively, these analyses demonstrate that *T12CS.7* encodes a previously uncharacterized B-type cyclin, now designated *cyb-4* (Cyclin B-4). *cyb-4* is predicted to encode three protein isoforms; although premature stop caused by *vv50* eliminates > 80% of the coding sequence of each, we confirmed that it was a null allele by generating a frameshift and premature stop codon immediately downstream of the initiating methionine using CRISPR-Cas9 genome editing. The engineered *cyb-4(vv177)* deletion suppressed the embryonic lethality and bivalent formation defects of *him-3(vvc)* mutants to the same extent as the *cyb-4* truncation (Fig. 1b and c), confirming that *cyb-4(vv50*) is a null allele and that the absence of CYB-4 mitigates *him-3(vvc)* meiotic defects.

The *him-3(vvc)* mutation results in a phenylalanine substitution at a highly conserved serine (S35) in the HORMA domain that impairs binding to HTP-3 and delays axis formation (Zetka et al. 2025). To understand the basis of *cyb-4(vv50)*-mediated suppression of *him-3(vvc)* mutant phenotypes we first assayed the HIM-3/HTP-3 colocalization coefficient (CoC) at meiotic entry as a measure of axis assembly (Fig. 1d and g). In the transition <u>z</u>one (TZ) of wild-type germlines (the region corresponding to leptotene-zygotene stages), chromatin is polarized within the nucleus and HTP-3 at the assembling axis recruits and colocalizes with HIM-3 (CoC = 0.872). In *him-3(vvc)* mutants, however, TZ nuclei are less polarized and HIM-3/HTP-3 colocalization is reduced to 60.6% of wild type (CoC = 0.529), consistent with previous reports of defective and delayed HIM-3 loading (Couteau et al. 2004; Zetka et al. 2025). In suppressed *him-3(vvc)* mutants, however, the colocalization of HTP-3 and HIM-3 rose to 73.7% of wild type (CoC = 0.643), indicating that loss of *cyb-4* partially restored timely axis formation (Fig. 1d and e). By midpachytene stages in wild-type nuclei, HIM-3 showed near-complete colocalization with HTP-3 along the chromosome axes (CoC = 0.901). Although HIM-3/HTP-3 colocalization remained modestly reduced at this stage in *him-3(vvc)* mutants in comparison to wild type (CoC = 0.874), it was fully restored to wild-type levels in *cyb-4(vv50); him-3(vvc)* suppressed mutants (CoC = 0.910) (Fig. 1f and g). Together, these results demonstrate that the early HIM-3 loading defect of *him-3(vvc)* mutants is partially suppressed by loss of *cyb-4*, and that once loaded, HIM-3 remains stably bound to HTP-3.

Because HIM-3 localization to chromosome axes is essential for SC formation (Couteau et al. 2004), we next asked if the improved axis morphogenesis in *cyb-4(vv50); him-3(vvc)* mutants was accompanied by improved SC assembly (Fig. 1h–k). In the TZ of wild-type germlines, SYP-1 colocalization with HTP-3 along chromosome axes is largely complete (CoC = 0.890). In *him-3(vvc)* mutants, however, the defect in HIM-3 localization results in extensive partial synapsis that reduced SYP-1/HTP-3 colocalization to 41.6% of wild type (CoC = 0.371), but improved to 62.2% in the presence of *cyb-4(vv50)* (CoC = 0.554) (Fig. 1h and i). By mid-pachytene, wild-type nuclei showed complete SYP-1/HTP-3 colocalization (CoC = 0.969). While colocalization levels in *him-3(vvc)* mutants reached 65.8% of wild type (CoC = 0.638), they rose to 91.7% in the presence of *cyb-4(vv50)* (CoC = 0.889) (Fig. 1j and k). These findings indicate that the improved HIM-3 axis localization that occurs in the absence of *cyb-4* is accompanied by substantial restoration of SC assembly.

In addition to its role in synapsis, HIM-3-mediated axis assembly is also required for the earlier recruitment of pairing center proteins to the <u>p</u>airing <u>c</u>enters (PC) (Bohr et al. 2016), sites located at one end of each chromosome that associate with the NE and are essential for homolog pairing (reviewed in Hillers et al. 2017). We next tested whether the improved HIM-3 localization to axes observed in suppressed *him-3(vvc)* mutants also restored PC assembly by monitoring the recruitment of PLK-2 (Polo-like kinase 2) to the PC binding proteins as a proxy (Fig. 1l and m) (Harper et al. 2011; Labella et al. 2011). In wild-type TZ nuclei, PLK-2 localization to PCs is visible as variably-sized aggregates at the nuclear periphery that represent transient clustering of the nuclear envelope-associated PCs during chromosome pairing stages (mean = 3.9 aggregates/nucleus). In *him-3(vvc)* mutants, most nuclei contained only a single PLK-2 aggregate corresponding to the X chromosome (mean = 1.0), consistent with the previously reported loss of autosomal homolog pairing (Couteau et al. 2004). The number of PLK-2 aggregates increased an average of 2-fold in *him-3(vvc)* suppressed mutants (mean = 2.0), indicating that *cyb-4(vv50)* partially restores PC formation and PLK-2 localization. Taken together, these results indicate that the loss of CYB-4 in *him-3(vvc)* mutants partially restores the levels of HIM-3 at chromosome axes, leading to concomitant improvements in HIM-3-dependent processes required for CO formation.

### CYB-4 is required for timely axis formation and SC assembly

To investigate the function of CYB-4 in meiosis, we first examined the localization of an endogenously tagged and fully functional FLAG::CYB-4 protein (Materials and Methods). CYB-4 is predominantly nuclear and localizes to nuclei throughout the germline, where it appears as nucleoplasmic puncta (Fig. 2). While both CYB-4 and CDK-1 show localization to germline nuclei (Brandt et al. 2020), the levels of CYB-4 sharply increase in TZ nuclei, consistent with the formation of a CYB-4/CDK-1 complex following meiotic entry.

**Fig. 2.**
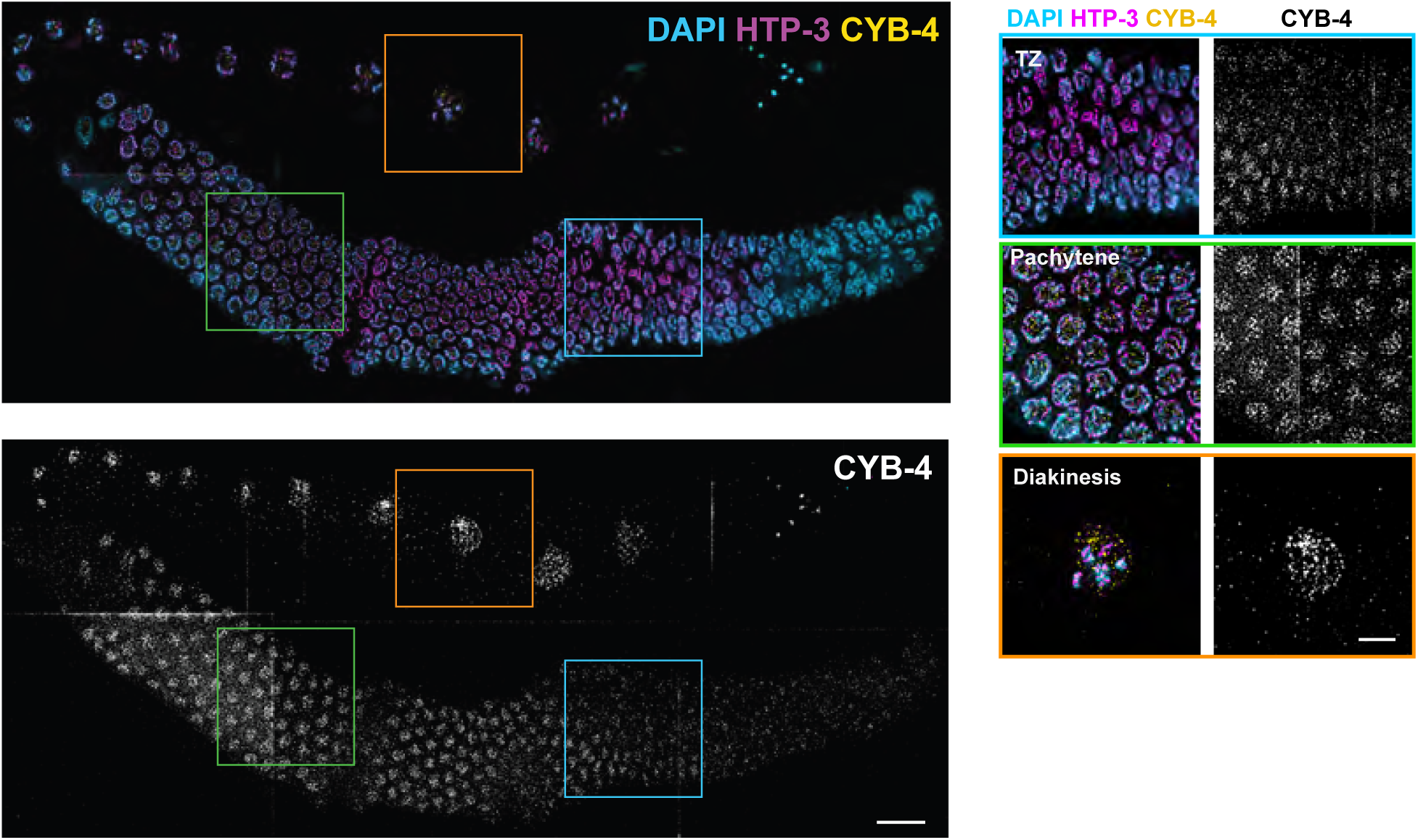
CYB-4 is highly enriched in meiotic nuclei. Immunofluorescence images of whole germlines stained for DNA, HTP-3 and CYB-4. Scale bar, 20µm. Magnified insets from the transition zone (TZ), midpachytene and diakinesis stages. Scale bar, 5µm.

*cyb-4(vv50)* and *cyb-4(vv177)* mutants consistently showed wild-type numbers of DAPI-stained bodies at diakinesis, indicating that any effect of CYB-4 loss on prophase events did not ultimately affect CO formation (Fig. 1c). Given that *cyb-4(vv50)*-mediated suppression of *him-3(vvc)* occurred during the earliest stages of axis assembly, we considered the possibility that the function of *cyb-4* was in regulating early prophase progression. To investigate this process, the meiotic entry region was defined by the mixing of nuclei positive for the meiosis-specific cohesin subunit COH-3/4 with nuclei lacking COH-3/4 staining (Severson and Meyer 2014). The length of the region (normalized to germline length) in *cyb-4(vv50)* mutants was increased 3-fold in comparison to wild type (Fig. 3a and b), indicating a delay in early prophase events. A hallmark of the mitotic-to-meiotic transition that occurs at the distal boundary of the TZ is the enrichment of HTP-3 on chromosomes as axis polymerization is initiated (Zetka et al. 2025). In *cyb-4(vv50)* mutants, HTP-3 localization in the TZ was delayed and disorganized; HTP-3/DAPI colocalization was reduced to ∼50% of wild type (CoC = 0.40 versus 0.76 in wild type), and HTP-3 and HIM-3 appeared as foci on the chromatin rather than as the contiguous stretches characteristic of wild-type nuclei at this stage (Fig. 3c and 3d). Furthermore, SYP-1/HTP-3 colocalization in TZ nuclei was similarly reduced in *cyb-4(vv50)* mutants (CoC = 0.49, CoC for wild-type = 0.89), and SYP-1 appeared in short stretches or variably-sized puncta, consistent with the delay in SC formation that occurs in the presence of axis assembly defects (Fig. 3e and f). By pachytene, however, HTP-3/DAPI colocalization was not different from wild type (P > 0.05) and chromosomes appeared fully synapsed (Fig. 3f), indicating that the early defect in axis assembly was resolved over time. Despite the formation of contiguous axes, however, the pachytene nuclei of *cyb-4(vv50)* mutants contained an uneven chromatin distribution with regions of aggregation and chromosomes that appeared longer (Fig. 3d), consistent with a broader defect in chromosome organization that is not restricted to axis assembly (Supplementary Fig. 3). Despite the delay in assembling chromosome structures and the later appearance of chromosome morphogenesis anomalies, the observation that *cyb-4(vv50)* mutants are competent for CO formation indicates that functional axes and SC are eventually formed, independent of CYB-4 function.

**Fig. 3.**
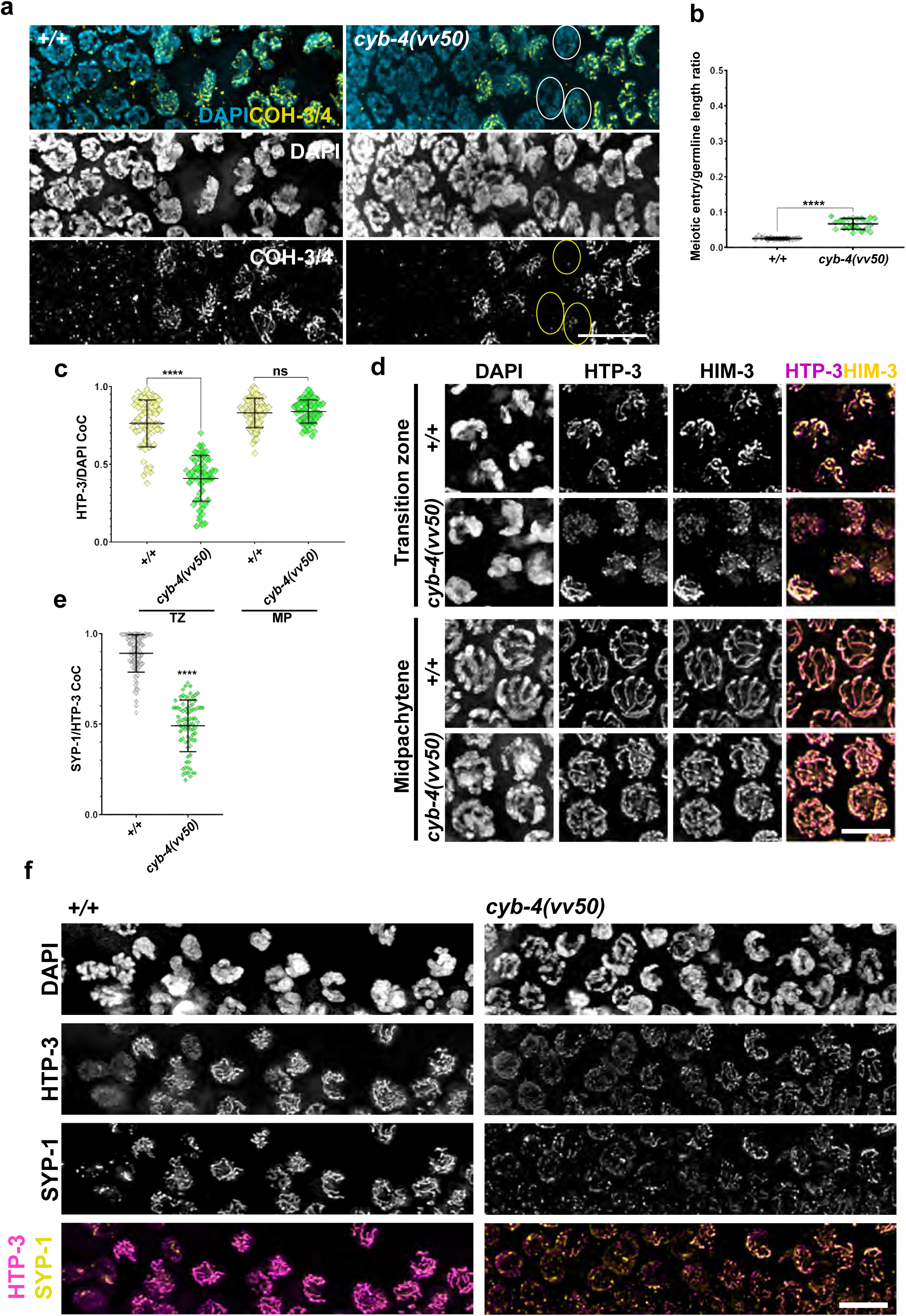
Meiotic entry and cell-cycle progression is delayed in *cyb-4(vv50)* mutants. a) Immunofluorescence micrographs showing stretches of the germline from the premeiotic region (left) to the TZ (right) stage stained for DNA and COH-3/4. Scale bar, 20µm. b) Scatterplot showing quantification of the ratio of the meiotic entry region length to the length of the entire germline in the indicated genotypes (n for wildtype = 15 and n for *cyb-4(vv50)* = 20). c) Scatterplot showing colocalization coefficient (CoC) of HTP-3/DAPI in the TZ and midpachytene of the indicated genotypes (n for TZ wildtype = 62, n for TZ *cyb-4(vv50)* = 56, n for MP wildtype = 66 and n for MP *cyb-4(vv50)* = 56). d) Immunofluorescence micrographs showing nuclei of the TZ and midpachytene stages stained for DNA, HTP-3 and HIM-3. Scale bar, 5µm. e) Scatterplot showing the CoC of SYP-1/HTP-3 in the TZ of the indicated genotypes. (n for wildtype = 96 and n for *cyb-4(vv50)* = 80). f) Immunofluorescence micrographs showing stretches of the germline from premeiotic region (left) to early pachytene (right) stained for DNA, HTP-3 and SYP-1. Scale bar, 5µm. Kruskal-Wallis test was used for statistical analysis and P < 0.05 was considered significant. ns: not significant, *: P < 0.1, **: P < 0.01, ***: P < 0.001, ****: P < 0.0001.

To investigate the effect of the delays in chromosome organization observed in *cyb-4* mutants on cell-cycle progression, we assessed the activity of the master meiotic regulator CHK-2. CHK-2 is activated at meiotic entry and remains active until midpachytene, however, in mutants that are defective in SC assembly and/or HR, its activity extends into later stages as part of meiotic checkpoint activation (MacQueen and Villeneuve 2001; Woglar et al. 2013; Kim et al. 2015; Zhang et al. 2023). In wild-type germlines, the inner nuclear envelope (NE) protein SUN-1 is phosphorylated at meiotic entry (Woglar et al. 2013); SUN-1 phosphorylation at S8 is dependent on CHK-2, and SUN-1^pS8^-specific antibodies were used to reveal CHK-2 activity and provide an insight into cell-cycle progression in *cyb-4(vv50)* mutants. In wild-type germlines, SUN-1^pS8^ is first found as numerous aggregates at the NE in TZ nuclei and then adopts a diffuse localization throughout the NE until mid-pachytene (Woglar et al. 2013). In *cyb-4(vv50)* mutants, however, SUN-1^pS8^ localization in the aggregates typical of TZ nuclei and in the NE was extended into late pachytene (1.6-fold increase in the CHK-2-active region relative to wild type), indicative of prolonged CHK-2 activity (Fig. 4a and b). The prolonged SUN-1^pS8^ signal in *cyb-4(vv50)* mutant germlines is consistent with the delayed loading of HIM-3 onto chromosome axes, given that defects in HIM-3 localization are known to extend CHK-2 activity (Kim et al. 2015). These results collectively indicate that CYB-4 regulates the events following meiotic entry and the tightly coupled processes that initiate chromosome morphogenesis and monitor the fidelity of axis assembly.

**Fig. 4.**
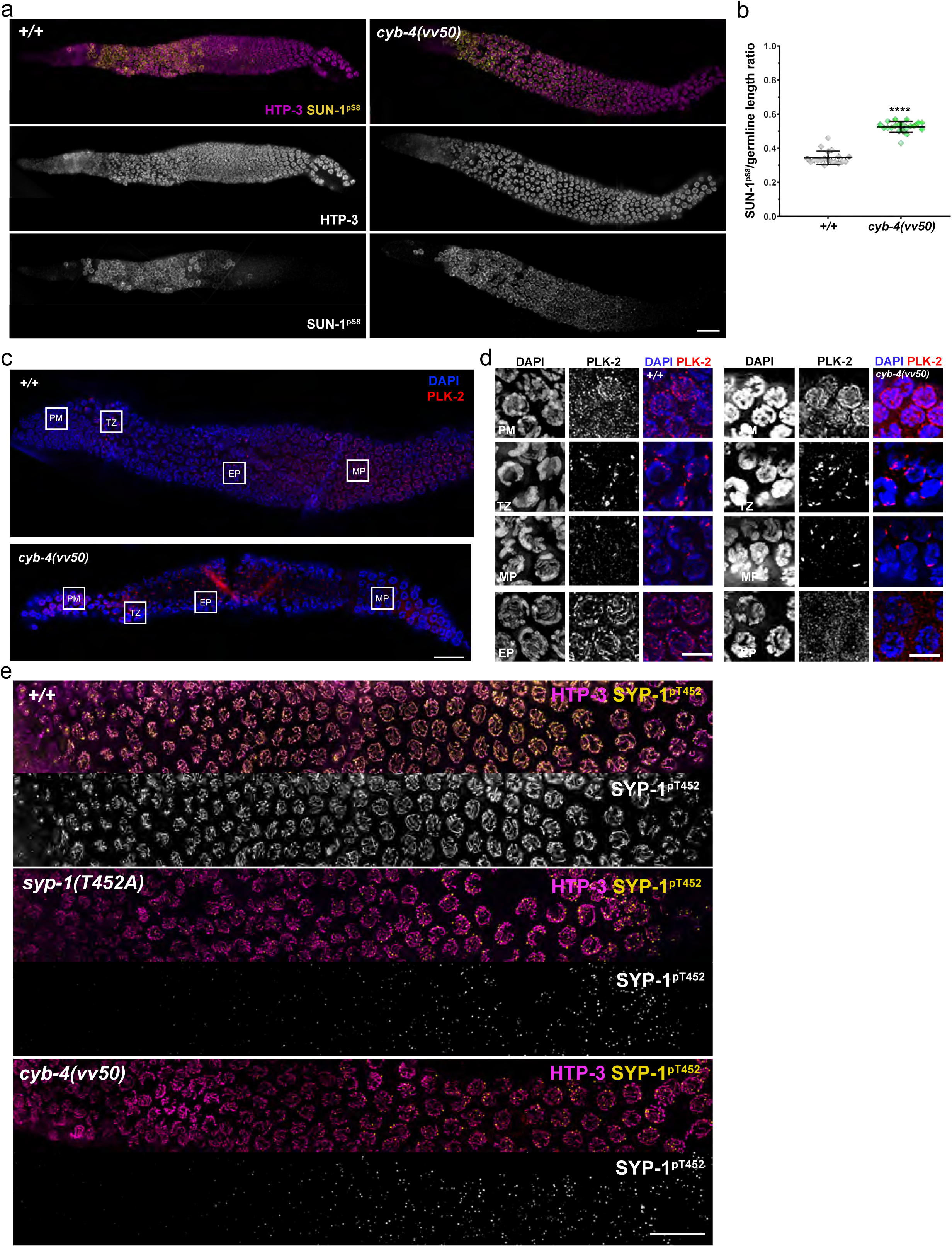
CYB-4 is required for SYP-1 phosphorylation and PLK-2 localization to the synaptonemal complex. a) Immunofluorescence images of the whole germlines from the distal tip (left) to the germline bend (right) stained for HTP-3 and SUN-1^pS8^. Scale bar, 20µm. b) Scatterplots showing quantification of active CHK-2 region over germline length (n for wild type = 19 and n for *cyb-4(vv50*) = 20). c) Immunofluorescence images of whole germlines from distal tip (left) to the germline bend (right) stained for DNA and PLK-2. Scale bar, 20µm. d) Immunofluorescence micrographs showing insets of premeiotic region (PM), transition zone (TZ), early pachytene (EP) and midpachytene (MP), from panel c. Scale bar, 5µm. e) Immunofluorescence images of the whole germlines from premeiotic region (left) to the late pachytene (right) stained for HTP-3 and SYP-1^pT452^. Scale bar, 20µm. Kruskal-Wallis test was used for statistical analysis and P < 0.05 was considered significant. ns: not significant, *: P < 0.1, **: P < 0.01, ***: P < 0.001, ****: P < 0.0001.

### CYB-4 is required for HR progression and timing of CO designation

Given the delay in axis assembly that was observed in *cyb-4(vv50)* mutants, we investigated if HR progression is also affected by measuring RAD-51 foci dynamics in a germline time-course analysis (Fig. 5a) (Colaiácovo et al. 2003). In wild type, RAD-51 foci first appeared in the TZ region, reached a peak in mid-pachytene, and were largely resolved by late pachytene. While RAD-51 foci appeared at wild-type-like levels and with appropriate timing in *cyb-4(vv50)* mutants they accumulated and persisted into inappropriately late stages, reaching an average 1.53 foci per nucleus in comparison to 0.11 foci per nucleus in wild type by late pachytene (Fig. 5b and c). These results indicate that while recombination initiation is not detectably affected in *cyb-4(vv50)* mutants, the chromosome morphogenesis defects that accompany the delayed entry into meiosis also delayed the timely repair of RAD-51-marked HR intermediates.

**Fig. 5.**
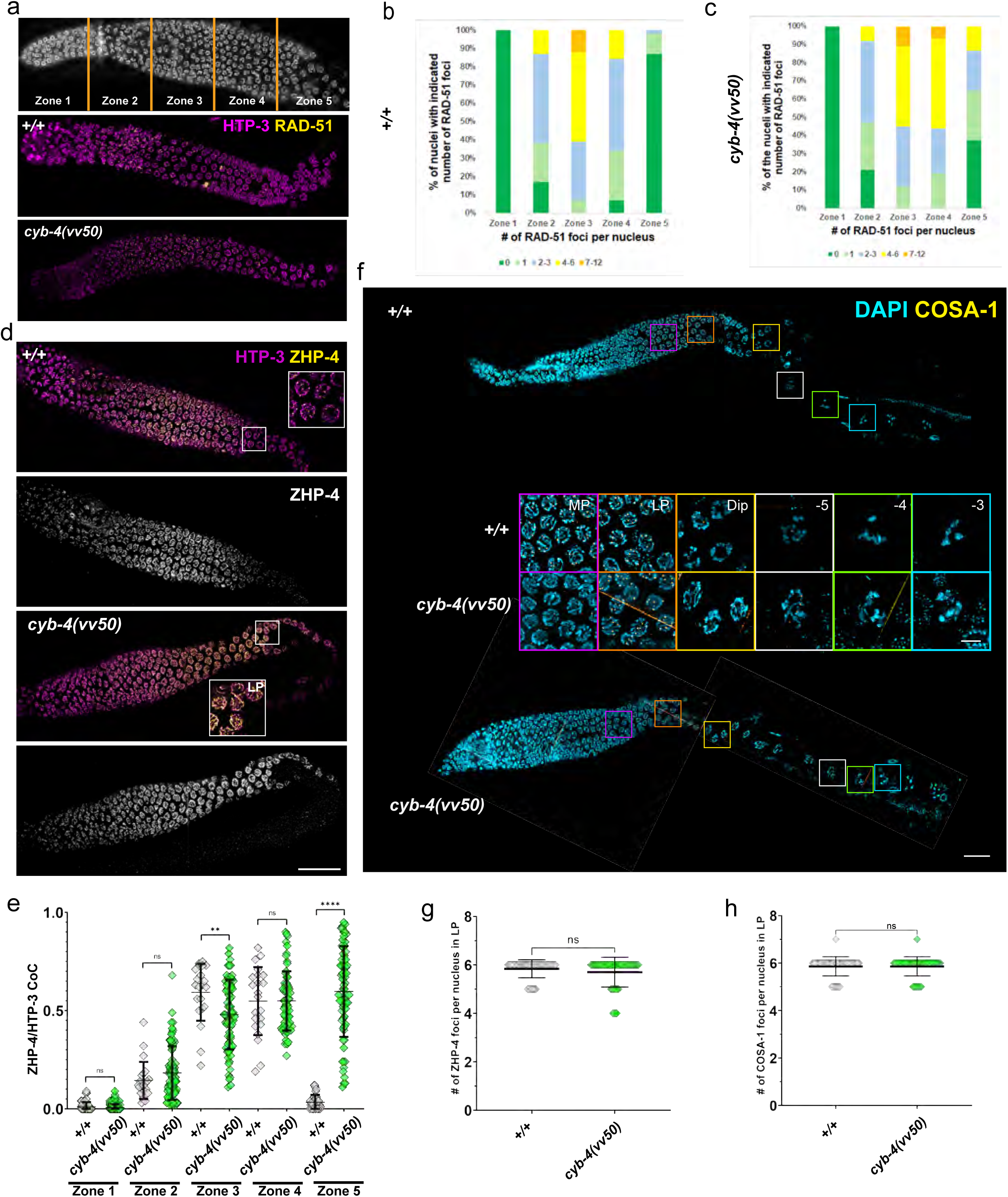
Homologous recombination is delayed in *cyb-4(vv50)* mutants. a) Germlines were divided into 5 equal zones from the distal tip to the proximal germline bend, roughly corresponding to mitotically proliferating region (Zone 1), TZ (Zone 2), and early, mid and late pachytene regions (Zones 3-5, respectively) and immunofluorescence images of the whole germlines from transition zone (left) to the germline bend (right) stained for HTP-3 and RAD-51. Scale bar, 20µm. b and c) Bar graphs showing time-course analysis of RAD-51 foci formation (a minimum of 100 nuclei were counted for each genotype in each zone). d) Immunofluorescence images of the whole germlines from TZ (left) to the germline bend (right) stained for HTP-3 and ZHP-4. Scale bar, 20µm. e) Scatterplots showing colocalization coefficient (CoC) of ZHP-4/HTP-3 in 5 zones of the germline. Germlines are divided into 5 equal zones from distal tip to the germline bend (a minimum of 40 nuclei were counted for each genotype in each zone). f) Immunofluorescence images of the whole germlines from distal tip (left) to the proximal (right) stained for DNA and COSA-1. Scale bar, 20µm. Insets are showing midpachytene (MP) in purple, late pachytene (LP) in orange, diplotene (Dip) in yellow, -5 diakinesis nuclei in white, -4 diakinesis nuclei in green and -3 diakinesis nuclei in blue. The scale bar of the insets, 5µm. g) Scatterplots showing the number of ZHP-4 foci per nuclei in late pachytene of the indicated genotypes. h) Scatterplots showing quantification of the number of COSA-1 per nucleus in late pachytene stage in the indicated genotypes. In g and h, a minimum of 50 nuclei were counted for each genotype. Kruskal-Wallis test was used for statistical analysis and P < 0.05 was considered significant. ns: not significant, *: P < 0.1, **: P < 0.01, ***: P < 0.001, ****: P < 0.0001.

To determine the effect of *cyb-4* loss on CO progression, we investigated the dynamics of proCO factors that localize to the 6 designated CO sites at late pachytene. In wild-type germlines, ZHP-4 first localizes along the SC during early-mid pachytene and becomes restricted to six foci that correspond to the designated CO sites by mid-to-late pachytene (Nguyen et al. 2018). In *cyb-4(vv50)* mutants, ZHP-4 appropriately localizes to the SC in early pachytene but persists along the SC into diplotene (Fig. 5d), indicating a delay in CO designation. To quantify these dynamics, germlines were divided into five equal zones from the distal tip to the gonad bend (Fig. 5a), and ZHP-4/HTP-3 colocalization was measured (Fig. 5e). In wild type, ZHP-4/HTP-3 colocalization first appears in the TZ, peaks in midpachytene, and abruptly drops as ZHP-4 is restricted to the designated CO sites. In *cyb-4(vv50)* mutants, the initial ZHP-4/HTP-3 CoC in the TZ is comparable to wild type (CoC in wild-type = 0.14 and in *cyb-4(vv50)* = 0.18), but ZHP-4 accumulates more gradually, peaks at late pachytene, and remains SC-associated until diplotene before restricting into six foci (Fig. 5g). These data indicate that both the recruitment to, and removal of, ZHP-4 from the SC are delayed in *cyb-4(vv50)* mutants; nevertheless, the eventual appearance of six ZHP-4 foci indicates that CO designation is intact despite delayed ZHP-4 kinetics. This conclusion is also supported by the localization dynamics of COSA-1, a marker of designated CO sites. In wild-type germlines, numerous small COSA-1 foci appear during mid-pachytene and then restrict to 6 large foci corresponding to the CO sites at mid-late pachytene (Fig. 5f and h) (Yokoo et al. 2012). In *cyb-4(vv50)* mutants, however, COSA-1-marked designated CO sites appear later (late pachytene) and persist longer, remaining detectable until the -3 diakinesis nucleus (Fig. 5f), indicating delayed CO designation and/or removal of proCO factors from CO sites. Despite this delay, the number of COSA-1 foci is not different from wild type (Fig. 5h), consistent with the presence of 6 bivalents at diakinesis. In summary, these results suggest that the meiotic entry delay observed in *cyb-4(vv50)* mutants results in downstream defects in chromosome axis assembly that trigger a compensatory extension of leptotene-zygotene to accommodate the CYB-4-independent processes of chromosome pairing, synapsis, and CO formation.

### CYB-4 is required for CDK-1-dependent meiotic prophase events

The role of CDK-1 in meiosis has been investigated using RNAi and auxin-inducible protein degradation, revealing that it is required for localization of PLK-2 to the SC and for bivalent remodeling at pachytene exit (Brandt et al. 2020). To evaluate if CYB-4 is the cyclin partner of CDK-1, we investigated the effect of CYB-4 loss on these processes.

An early role of CHK-2 following meiotic entry is the phosphorylation of PC-binding proteins to provide a docking site for PLK-2 binding during chromosome pairing stages (Kim et al. 2015); following CHK-2 inactivation at midpachytene, PLK-2 relocalizes from the PCs to the SC (Harper et al. 2011; Labella et al. 2011; Kim et al. 2015). Since SC localization of PLK-2 is dependent on phosphorylation of SYP-1 at T452 by CDK-1 (Sato-Carlton et al. 2018; Brandt et al. 2020), we investigated the dynamics of PLK-2 localization as a means of testing the model that *cyb-4* is required for CDK-1 activity. Consistent with the prolonged CHK-2 activity observed in *cyb-4(vv50)* mutants, PLK-2 persisted at PCs into regions corresponding to midpachytene in wild-type germlines (Fig. 4c and d). Importantly, PLK-2 failed to relocalize to the SC in midpachytene nuclei and instead exhibited a diffuse nuclear localization pattern into late pachytene stages that resembled the effect of CDK-1 loss (Brandt et al. 2020) (Fig. 4c and d).

We next investigated if the SYP-1^pT452^ phosphoepitope was lost in *cyb-4(vv50)* mutants using an antibody specific for phosphorylated SYP-1 (SYP-1^pT452^) (Sato-Carlton et al. 2018; Brandt et al. 2020). In wild-type germlines, SYP-1^pT452^ is first detected at PCs in the TZ, extends along the SC during pachytene, and subsequently becomes restricted to the short arms of bivalents at pachytene exit stages (Fig. 4e) (Nabeshima et al. 2005; Martinez-Perez et al. 2008). In contrast, *cyb-4(vv50)* mutants showed dramatically reduced SYP-1^pT452^ levels, comparable to those observed in the phosphodead *syp-1(T452A)* mutant (Fig. 4e). We considered the possibility that SYP-1 phosphorylation alone is sufficient for suppression of *him-3(vvc),* however embryonic lethality and CO defects were not suppressed in *him-3(vvc)*; *syp-1(T452A)* mutants (Supplementary Fig. 2), indicating that loss of SYP-1 phosphorylation at T452 is not the basis of the *cyb-4(vv50)* suppression. Taken together, these results indicate that CYB-4 is required for CDK-1-dependent phosphorylation of SYP-1 and provides strong evidence that it is the cyclin partner of CDK-1 during meiosis.

In addition to regulating PLK-2 relocalization through SYP-1 phosphorylation, CDK-1 also promotes chromosome condensation following pachytene exit and oocyte maturation (Brandt et al. 2020; Wenda et al. 2021). To determine if CYB-4 is also required for these later CDK-1-dependent events, chromosome morphology was first examined in diplotene and diakinesis nuclei using DAPI and HTP-3 staining. In wild type, pachytene exit correlates with progressive condensation of the chromatin and shortening of the desynapsed chromosome axes until highly compacted bivalents appear at diakinesis (Fig. 5a). Following pachytene exit in *cyb-4(vv50)* mutants, chromosome condensation was markedly delayed, with decompacted chromosomes persisting through the -3 and -2 diakinesis oocytes (-1 is the most proximal oocyte and nearest to the spermatheca, Fig. 5a), resembling previously reported CDK-1 depletion phenotypes (Brandt et al. 2020). In the -1 oocyte, however, six condensed bivalents were observed, indicating that chromosome condensation is delayed rather than abolished.

Given that loss of CYB-4 mimicked the CDK-1-depletion defects in chromosome compaction following pachytene exit, we investigated if the *cyb-4(vv50)* mutants also displayed defects in the chromosome remodeling that accompanies SC disassembly at the late pachytene-diplotene transition. During this process, CO formation triggers partitioning of the bivalent into discrete long and short arms, which can be visualized by the differential localization of chromosome markers (Martinez-Perez *et al*. 2008; de Carvalho *et al*. 2008). In wild-type germlines, SYP-1 and ZHP-2 are lost from the long arms of the bivalent and retained on the short arms until the -3 oocyte, while the axis component HTP-1 shows the reciprocal pattern and is retained on the long arms (Fig. 6). Despite the defects in chromosome condensation observed in *cyb-4(vv50)* mutants, SYP-1, ZHP-2, and HTP-1 showed reciprocal localization along the bivalent, indicating that arm partitioning was established (Fig. 6); nevertheless, bivalent formation was delayed, with fully formed and condensed bivalents appearing only at the -1 diakinesis nucleus. Together these results demonstrate that CYB-4 is not required for CO designation, but is required for the timing of CO-triggered chromosome remodeling.

**Fig. 6.**
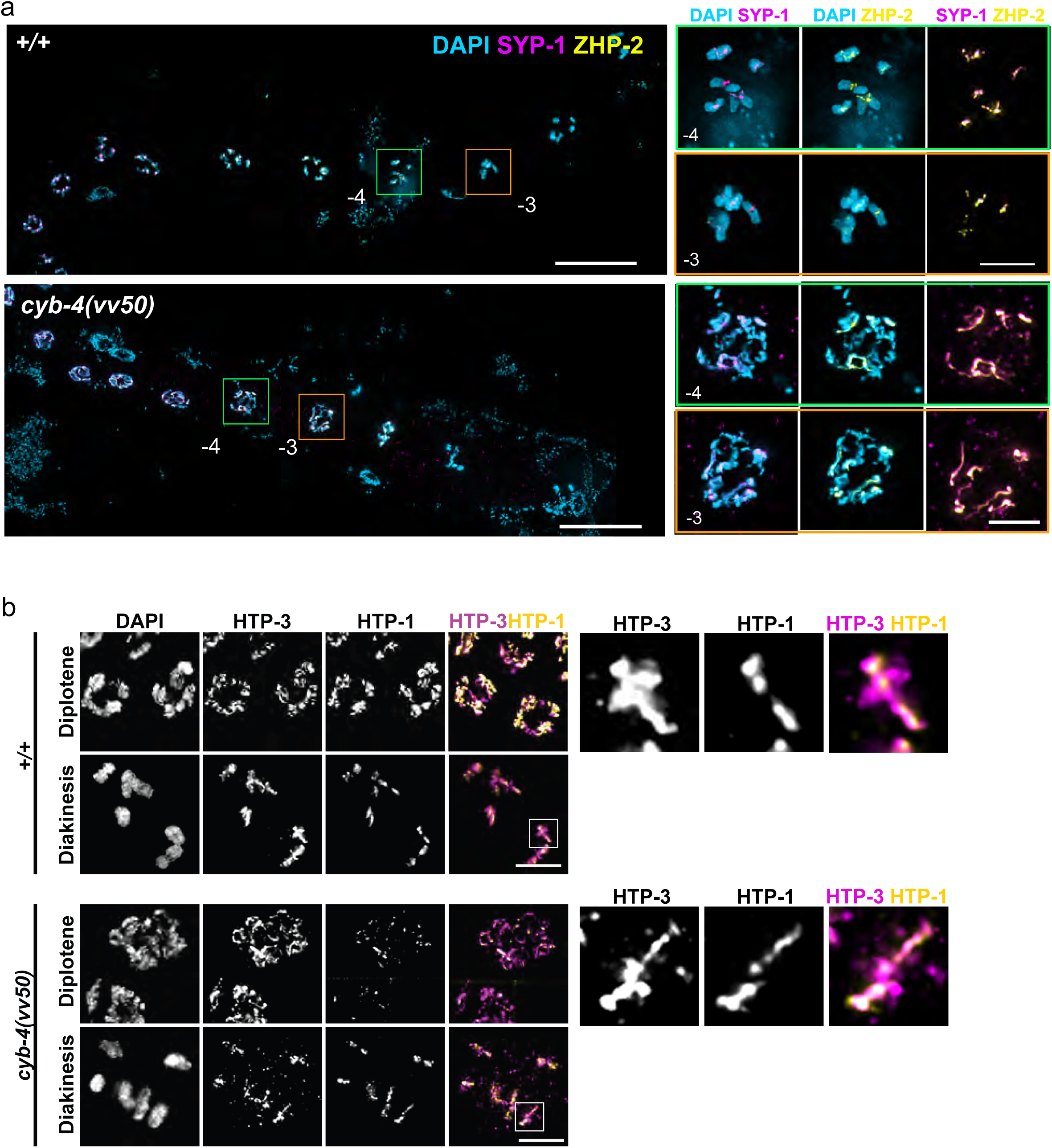
CYB-4 is required for the timing of chromosome remodeling. a) Immunofluorescence images of the germline stretch from diplotene (left) to the diakinesis stage stained for DNA, SYP-1 and ZHP-2. Insets show magnified regions of -4 (green) and -3 (orange) diakinesis nuclei. b) Immunofluorescence micrographs of the diplotene and diakinesis stages stained for HTP-3 and HTP-1. Insets show magnification of single bivalents. Scale bar, 20µm. Inset scale bars, 5µm.

Since CDK-1 is also required for oocyte maturation, we used the appearance of phosphorylated histone H3 (pH3) as a marker to investigate the process (Hsu *et al*. 2000; reviewed in Kim *et al*. 2013). In wild-type germlines, pH3 first appears on the short arms of bivalents at the -3 oocyte and progressively accumulated through the -1 oocyte (Fig. 7a). In contrast, *cyb-4(vv50)* mutants exhibited a pronounced delay in pH3 accumulation, with signal detected only in the -1 oocyte (Fig. 7b), closely phenocopying the effect of CDK-1 depletion (Hsu *et al*. 2000; Ferrandiz *et al*. 2018). Collectively these results demonstrate that *cyb-4(vv50)* mutants exhibit multiple defects characteristic of depleted CDK-1 activity, including prolonged PLK-2 retention at pairing centers, drastically reduced SYP-1^pT452^ phosphorylation, and delays in chromosome compaction, bivalent remodeling, and oocyte maturation. These findings support the conclusion that CYB-4 is the meiotic cyclin that promotes CDK-1 activity during meiotic prophase and oocyte development.

**Fig. 7.**
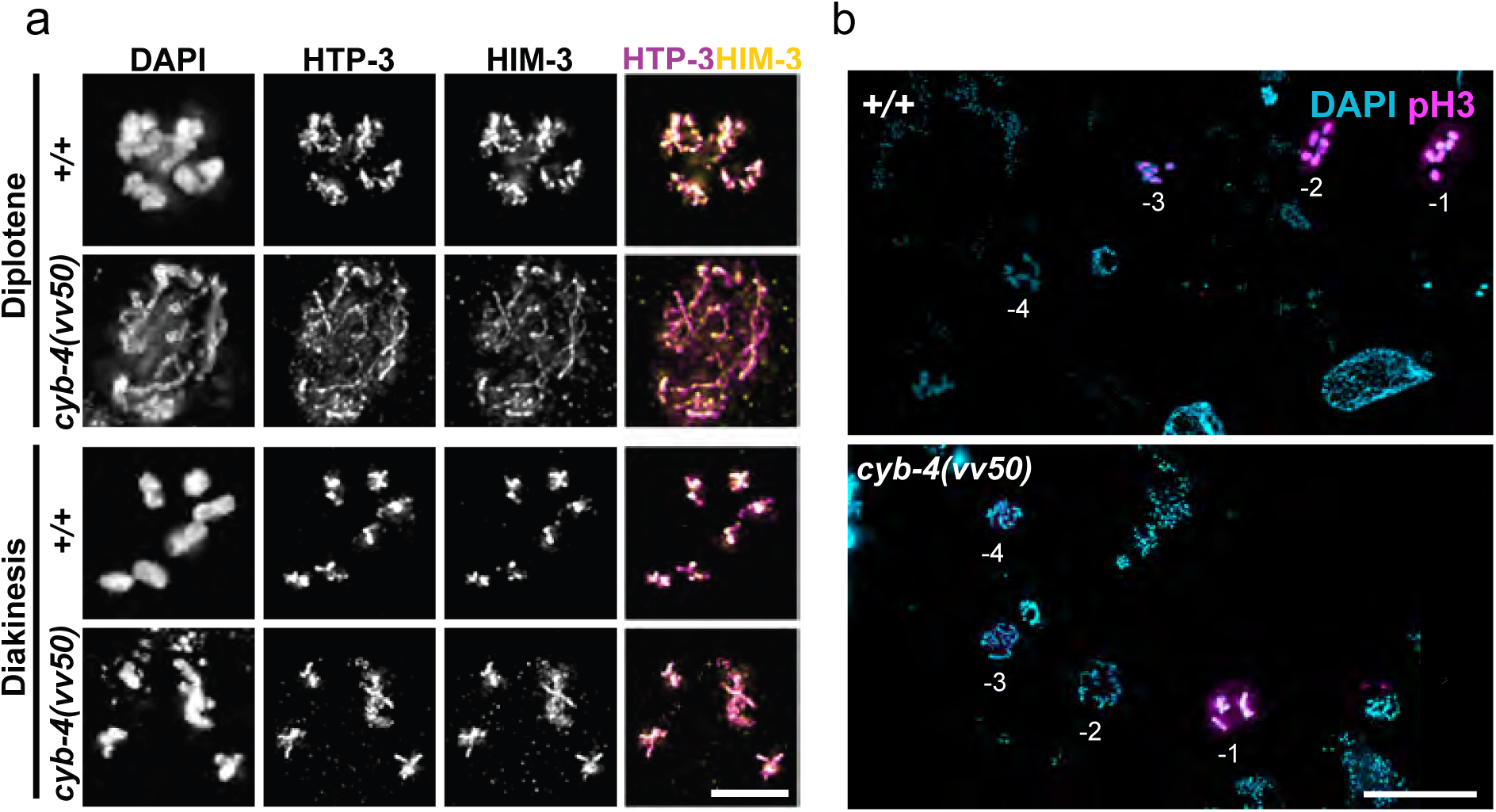
CYB-4 is required for chromosome condensation at late meiotic prophase and timely oocyte maturation. a) Immunofluorescence micrographs showing nuclei of diplotene and diakinesis stages stained for DNA, HTP-3 and HIM-3. Scale bar, 5µm. b) Immunofluorescence images showing germline stretches from diplotene to diakinesis stages stained for DNA and phospho-histone-3 (pH3). Scale bar, 20µm.

## Discussion

Successful meiosis depends not only on the completion of chromosome-associated events but also on their precise temporal coordination with cell-cycle progression. Here, we reveal CYB-4 to be a previously uncharacterized meiosis-specific B-type cyclin that is required for the correct timing of axis assembly, bivalent remodeling, and oocyte maturation. Our results are compatible with the model that CYB-4 is the cyclin partner of CDK-1, and that it directs CDK-1 activity to meiotic prophase targets regulating these events. Consistent with a central role for CYB-4/ CDK-1 complex, *cyb-4(vv50)* mutants exhibited phenotypes that mimic those previously described in CDK-1-depleted germlines: a loss of SYP-1^pT452^ phosphorylation together with defective recruitment of PLK-2 to the SC (Sato-Carlton et al. 2018; Brandt et al. 2020), defects in bivalent decondensation and remodeling, and pH3 accumulation during oocyte maturation (Hsu *et al*. 2000; Ferrandiz *et al*. 2018). The phenotypic parallels between *cyb-4* and *cdk-1* loss strongly suggest that the proteins function in the same pathways, consistent with the formation of a CYB-4/CDK-1 complex. While the requirement for CDK-1 in post-CO remodeling has been previously reported (Brandt et al. 2020), our results expose an earlier function following meiotic entry that has been experimentally inaccessible using RNAi or auxin-inducible degradation of CDK-1 given the confounding cell-cycle defects that accompany CDK-1 loss in the proliferating germline. Our findings support a model in which CYB-4 partners with CDK-1 during meiotic prophase to define windows of chromosome morphogenesis to create switch-type transitions at meiotic prophase entry and pachytene exit.

Our results indicate that *cyb-4* is not required for meiotic entry, but is instead required for the timely execution of prophase events. Chromosome axis formation is one of the earliest visible hallmarks of meiotic entry, and *cyb-4(vv50)* mutants show delayed assembly of axis components that is accompanied by delayed synapsis and CO designation. These interdependent processes are nevertheless eventually completed by late pachytene, indicating that the early delay is absorbed during pachytene progression, leaving CO formation unaffected. Following CO designation at late pachytene, *cyb-4(vv50)* mutants also exhibit a delay in pachytene exit which postpones bivalent remodeling, chromosome compaction, and oocyte differentiation; these delays are also overcome to eventually produce cytologically normal diakinesis oocytes that support wild-type-like levels of embryonic viability. Together these observations indicate that the primary role of CYB-4 is not in determining if individual meiotic processes occur, but in regulating the timing of two major cell-cycle transitions at which chromosome organization changes dramatically: one at meiotic prophase entry and a second at pachytene exit. Our results suggest that following meiotic entry, CYB-4 is required to constrain axis assembly to a defined window; its extension in *cyb-4* mutants results in slower axis assembly that in turn causes SC defects and sustained CHK-2 activity. At the second step, CYB-4 defines the window corresponding to pachytene exit and its expansion results in a temporal shift in bivalent remodeling, chromosome compaction, and oocyte maturation. Our results are compatible with a model in which CYB-4 is the cyclin partner of CDK-1 during meiotic prophase, directing CDK-1 activity toward prophase-specific substrates to time switch-like transitions at meiotic prophase entry and pachytene exit. These are points at which chromosome organization changes irreversibly, and consequently the timing of prophase events is as critical as their completion.

A surprising element of this study is the finding that loss of *cyb-4* suppresses multiple meiotic defects associated with expression of the binding-defective HIM-3^S35F^ protein. Since it is improbable that loss of CYB-4 could restore the binding capacity of the mutant HIM-3 protein, we favor the model that expansion of the window of axis assembly that occurs in the absence of CYB-4 provides additional time for the loading-defective HIM-3^S35F^ protein to bind to HTP-3, a proposition that is supported by the partial restoration of HIM-3 loading in *him-3(vvc); cyb-4(vv50)* mutants, and the accompanying improvements in homolog pairing and CO formation.

In conclusion, we identify CYB-4 as a previously uncharacterized, meiosis-specific B-type cyclin that directs CDK-1 activity during meiotic prophase. Rather than being required for individual meiotic events to occur, CYB-4 controls their timing by defining two windows in which chromosome organization undergoes irreversible change: at meiotic prophase entry to promote timely chromosome axis assembly, and at pachytene exit to regulate bivalent restructuring and oocyte differentiation. CYB-4 thus specializes CDK-1 activity to establish an additional layer of temporal control to chromosome morphogenesis that ensures that structural intermediates do not linger and interfere with tightly coordinated prophase processes.

## Acknowledgements

The authors would like to thank the members of the Zetka lab for discussions and critical reading of the manuscript. Antibodies, strains, and other reagents were generously provided by Yumi Kim (Johns Hopkins University), Nicola Silva (Masaryk University), Monica Colaiácovo (Harvard University), and Verena Jantsch (University of Vienna).

